# The Limits of Touch: Spatial acuity for frequency-resolved air-borne ultrasound vibrotactile stimuli

**DOI:** 10.1101/2023.02.08.527028

**Authors:** Antonio Cataldo, William Frier, Patrick Haggard

## Abstract

Spatial acuity is a fundamental property of any sensory system. In the case of the somatosensory system, the two-point discrimination (2PD) test has long been used to investigate the spatial resolution of tactile perception. The somatosensory system comprises multiple mechanoreceptive channels, each tuned to specific vibrotactile frequencies. In particular, the rapidly adapting channel (RA) responds to low-frequency vibration and is thought to have high spatial acuity. The Pacinian channel (PC) responds to high-frequency vibration and is thought to convey little or no spatial information. However, the mechanical stimulations used in most 2PD tests make it difficult to disentangle the contribution of each mechanoreceptive channel to spatial tactile perception. Here we developed a novel 2PD test based on ultrasound stimulation to deliver frequency-resolved vibrotactile stimuli designed to preferentially activate specific tactile channels. Across four experiments, we systematically investigated the spatial resolution of the two main vibrotactile channels. Contrary to the textbook view of poor spatial resolution for PC-like stimuli, we found that high-frequency vibration produced surprisingly good spatial acuity. This effect remained after controlling for differences between the channels in stimulus detectability and perceived intensity. Laser doppler vibrometry experiments confirmed that the acuity of the PC channel was not simply an artifact of the skin’s resonance to high-frequency mechanical stimulation. Thus, PC receptors may transmit substantial spatial information, despite their sparse distribution, deep location, and large receptive fields.

## Introduction

A fundamental property of any sensory system is its spatial resolution or acuity, defined as the minimal distance required by a given sensory system to distinguish between two adjacent stimuli. Several studies have investigated spatial acuity for visual^1^ and auditory stimuli^2^. However, the case of somatosensation is more complex, because the somatosensory system comprises at least three main mechanoreceptive channels, each preferentially responding to different types of skin deformation, and having different distributions of spatial receptors across the skin^3^. Slowly adapting type 1 (SAI) afferents terminate in Merkel disks and are most sensitive to static pressure and low frequencies (<10 Hz). Rapidly adapting (RA) afferents innervate Meissner corpuscles and are preferentially activated by flutter-like vibrations (10 – 50 Hz). Finally, Pacinian afferents (PC) innervate Pacinian corpuscles and are sensitive to high-frequency vibrations (> 50Hz, peaking at 250 Hz).

Tactile spatial acuity is traditionally measured using the two-point discrimination (2PD) test (see for example^4^). In this task, a calliper-like object is used to apply pressure on two adjacent points on the skin of the observer, which is asked to report whether they feel one or two stimuli. The distance between the two points is systematically manipulated in successive trials so to establish the minimal distance at which the observer can correctly distinguish two separate points from a single point presented at the central position. Tactile acuity has been extensively studied both in clinical^5–7^ and experimental^8–12^ settings (for a review see^13^). The two-point approach has been criticised^14^, and improved measurement methods have been proposed^15^, though these are only applicable to some skin regions. Essentially, the 2PDT remains a key method for assessing tactile sensory function, and the general approach has remained substantially unchanged since its first mention in Weber’s foundational work^16,17^.

In particular, the overwhelming majority of previous works used mechanical static-pressure stimuli to measure 2PD thresholds. This clearly focusses study of tactile acuity on the SA-I mechanoreceptor channel alone, and largely ignores other mechanoreceptive channels. In particular, it is still unclear what the contribution of the frequency-specific RA and PC mechanoreceptive channels to spatial perception might be, and how stimulus features such as amplitude and frequency affect the spatial resolution capabilities of the somatosensory system.

Several studies since Weinstein’s seminal work^4^ have advanced the hypothesis that the tactile spatial acuity of a specific body area is determined by two main factors: the size and the innervation density of the receptive field^12,18^. For example, whole-body mapping studies have revealed gradients of tactile spatial acuity consistent with the innervation density of the mechanoreceptors^12^. Interestingly, classical physiological evidence shows that the relative innervation density of RA afferents on the palm and the intermediate/proximal phalanges is, respectively, 2.5 and 4 times higher than the innervation density of PC afferents^19^. Thus, an account based on the idea that 2PD thresholds depend only on the innervation density would predict that stimuli preferentially activating the RA afferents should show a much greater spatial acuity than stimuli preferentially activating the PC afferents. Indeed, many textbooks state that due to the extreme sensitivity, low density, high depth, and large size of their RF, PC afferents convey little or no spatial information^3^. However, this hypothesis has not been directly tested yet.

Some previous studies attempted to address this knowledge gap with specialized stimulators allowing multipoint stimulation with variable distances and vibrotactile frequencies^20–23^. However, the design of those stimulators means that a constant indentation (i.e., static pressure) must be applied to the participant’s skin to deliver the required vibrotactile patterns. For this reason, tactile stimuli at any frequency are typically superimposed upon a background of steady pressure. Therefore, the results reported by those studies may be influenced by mechanical interactions between propagation waves on the skin^24,25^ or the various neural interactions between the SA1 and the RA/PC channels^9,26,27^. An alternative approach might involve positioning a mechanical probe so that it just contacts the skin, without any steady pressure. However, if the probe is then vibrated sinusoidally, it can only give a half-cycle of stimulation, and must then break contact for the other half of the cycle, resulting in no stimulation at all. For these reasons, estimates of the spatial acuity of individual frequency-resolved mechanoreceptor channels (RA, PC) remain elusive, and the spatial limits of human touch submodalities are not well-known.

Here, we addressed this question using a novel, contactless type of tactile stimulation that produces precise frequency-resolved vibrotactile patterns without the steady pressure or static indentation that accompany mechanical contact. We used focussed ultrasound stimulation to project discrete points of acoustic pressure, thus generating contactless tactile sensations^28^. Crucially, since ultrasound stimulation does not require any mechanical stimulator to touch the skin, coactivation of SA receptors is avoided, in contrast to classical mechanical vibrotactile stimulation. Moore and colleagues^29^ used microneurography to confirm that median nerve units classified as low-threshold, SAI and SAII nerves, did not respond to ultrasound stimuli delivered to their receptive field by a device similar to the one used in the present study.

Across four experiments, we systematically investigated the spatial acuity of multiple somatosensory mechanoreceptor channels using pure frequency-resolved stimuli. In Experiment 1, we compared 2PD thresholds for low- (50 Hz) and high-frequency (200 Hz) ultrasound stimuli delivered either on the palm or the index finger of their left hand. In Experiment 2, we investigated whether frequency-specific differences in spatial acuity could be secondary consequences of frequency-specific differences in perceived stimulus intensity, by adjusting the physical intensity of different frequency-specific stimuli so that they were matched in terms of *perceived* intensity. Experiment 3 provided a preregistered replication of the results found in the previous experiments and aimed to establish whether effects of stimulus frequency and perceived intensity on spatial acuity were mediated by stimulus detectability. Finally, in Control Experiment 4, we used a laser doppler vibrometer (LDV) to measure the physical peak-to-peak displacement induced by our ultrasound stimuli on the skin. This allowed us to disentangle whether differences in perceived intensity and spatial acuity for different stimulus frequencies were due to frequency-specific mechanical resonances in the skin^24,25^, or to the selective tuning of the different neural receptors.

## Results

### Does the frequency of vibrotactile stimuli affect tactile spatial acuity?

Experiment 1 aimed to test whether the frequency and skin region of a vibrotactile stimulus affected the participants’ 2PD threshold. In a 2 (Frequency: 50, 200 Hz) x 2 (Skin Region: palm, index finger) within-participants design, participants (n = 12) performed a classical 2PD task based on adaptive staircases^12^ (see Methods and Figure S1). We used a novel ultrasound tactile stimulation that allows the projection of tactile points directly onto the skin of the participants^28^. Crucially, the location and amplitude of the focal point generated by the device can be spatiotemporally modulated to produce vibrotactile patterns of different shapes and frequencies^30^. Thus, this contactless stimulation allowed us to delivered pure 50 Hz and 200Hz frequency-resolved stimuli with minimal activation of the pressure-sensitive SAI and SAII afferents^29^. Participants received single (~20% of the trials) or two-point (~80% of the trials) stimuli on the palm of their hand or on their index finger and were asked to report whether they felt a single or a double point (see Methods). The distance between the two points was adaptively adjusted until a threshold value was reached where participants could no longer discriminate between one- and two-point stimuli (see Methods).

Classical physiological evidence suggest that the relative innervation density of RA afferents on the palm and the intermediate/proximal phalanges is, respectively, 2.5 and 4 times higher than the innervation density of PC afferents^19,31,32^. Based on this physiological evidence, we originally hypothesised a significant main effect of the frequency factor (see preregistration at https://osf.io/jkd8h), with the 50 Hz stimuli (preferentially activating the RA channel) producing a significantly lower 2PD threshold (i.e., higher acuity) than the 200 Hz stimuli (PC channel). We also expected a significant interaction driven by a significantly lower 2PD threshold for the 50 Hz stimuli on the index finger (which contains relatively more RA afferents)^19^, compared to the palm of the hand.

Surprisingly, and contrary to our preregistered hypothesis, we found a significant main effect of Frequency (*F_1,11_* = 7.26; *p* = .021; _η_ *_p_* = .398) showing that the participants’ 2PD thresholds was in fact *lower* (15.62 ± 7.06 mm) for the 200 Hz stimuli (preferentially activating the sparser PC afferents) than for the 50 Hz stimuli (21.80 ± 6.44 mm) (preferentially activating the more ubiquitous RA afferents). Furthermore, the main effect of Skin region (*F_1,11_* = 0.47; *p* = .509; _η_ *_p_* = .041; B_01_ = 2.87, error % = .021) and the interaction between factors were both non-significant (*F_1,11_* = 1.89; *p* = .196; _η_ *_p_* = .147; B_01_ = 1.62, error % = .005), suggesting that the effect of frequency on spatial acuity was not dependent on the stimulation skin region.

**Figure 1.**
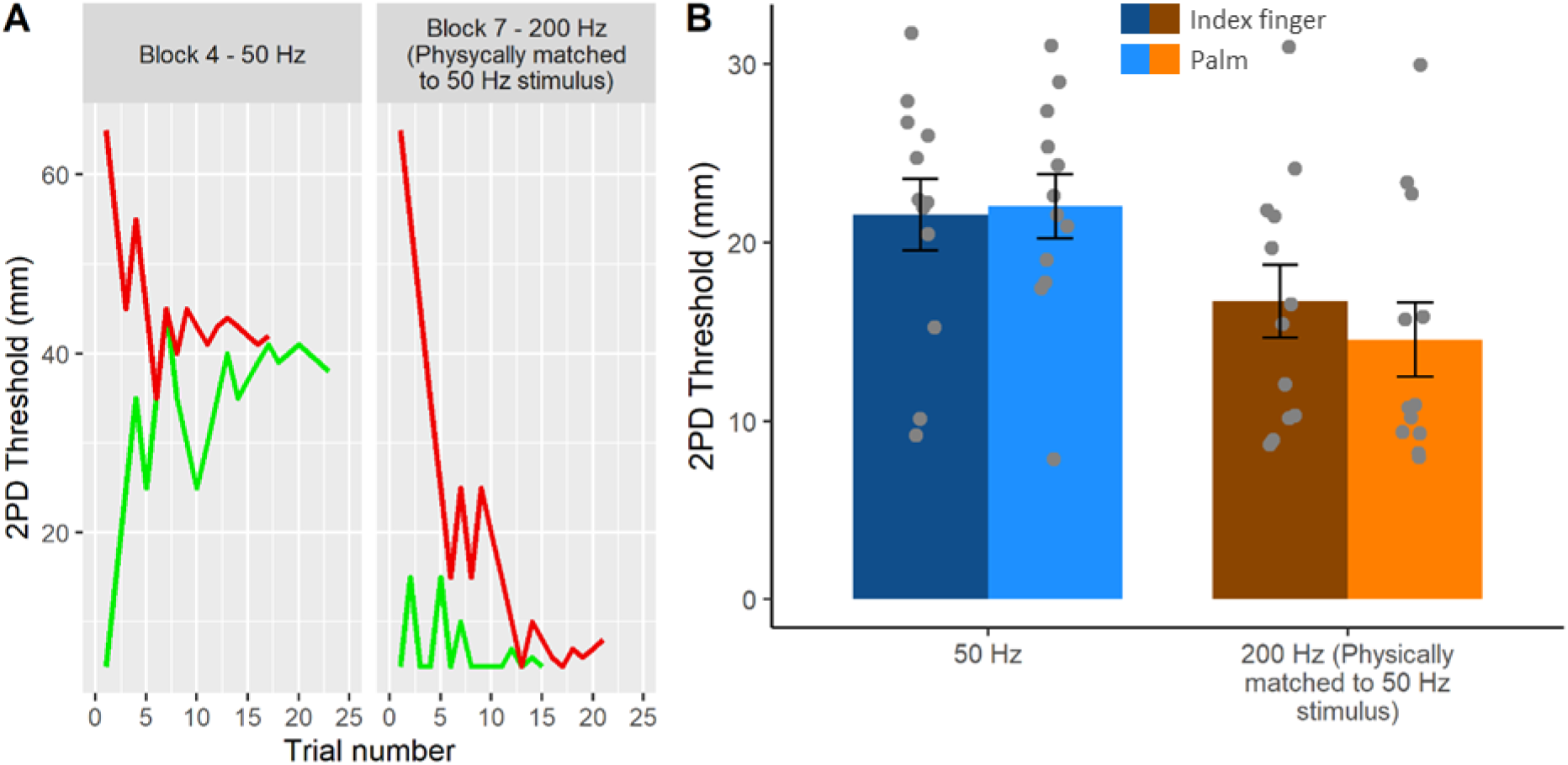
Results from Experiments 1. **A. Staircase data from the 50 Hz and physically matched 200 Hz conditions in a representative participant (P#2, blocks 4 and 7).** We used a staircase procedure to identify the smallest distance at which two points are no longer discriminable from a single centrally-located point. Red an green lines show responses to descending and ascending staircases, respectively. **B. Two-point discrimination thresholds.** The staircase procedures yield estimates of two-point discrimination threshold (2PDT), for 50 Hz an for200 Hz, on the finger (darker colours) and on the palm (lighter colours). A lower 2PDT corresponds to better spatial resolution. Error bars represent the SEM.

### Does the frequency of vibrotactile stimuli affect their perceived intensity?

The ultrasound stimuli used in Experiment 1 were matched in terms of physical intensity (see Methods). However, verbal reports from the participants in the debriefing session at the end of the experiment suggested that the perceived intensity of the 200 Hz stimulus might be higher than that of the physically matched 50 Hz stimulus. The high sensitivity of the PC is well-established, with microneurographic studies showing that skin indentations as small as 10 nm are sufficient to trigger an action potential in afferents associated with PCs^3,31,33^. In contrast, the sensitivity of the RA afferents is about two orders of magnitude lower^3^. To investigate whether the counterintuitive results found in Experiment 1 were due to the difference in the perceived intensity of the stimuli, Experiment 2 began by participants matching the intensity of 50 Hz and 200 Hz stimuli. The intensity matching task was based on a 2-AFC staircase procedure like the one described above for the 2PD thresholds (see Methods). Participants received a (reference) 50 Hz stimulus and a 200 Hz (comparison) stimulus in each trial and were asked to report which of the two stimuli felt stronger.

A direct comparison between the participants’ average PSE for 200 Hz stimuli against the test value of 1 (i.e., the intensity of the 50 Hz stimulus) showed that for equal stimulus energy, the 50 Hz stimuli felt significantly less intense than the 200 Hz stimuli (70.92 ± 14.41 %; t_11_ = −6.99; *p* < .001; d_z_ = −2.018).

### Does the perceived intensity of the stimuli mediate the effect of frequency on spatial acuity?

In the second part of Experiment 2, we used the same 2PD procedure implemented in Experiment 1 to test the spatial acuity of 50 and 200 Hz stimuli that were equal in perceived intensity. Given that the Skin region factor in Experiment 1 was not significant, we restricted testing to the palm alone. Thus, Experiment 2 had a single factor (Frequency) with two levels (50, 200 Hz).

If the high spatial resolution for 200 Hz compared to 50 Hz stimuli observed in Experiment 1 was due to the difference in the perceived intensity of the stimuli, then adjusting the two stimuli to equalise perceived intensity should remove the apparent 200 Hz advantage for spatial resolution (hypothesis 1), or even reverse it (hypothesis 2). Conversely, if the effect found in Experiment 1 did not depend on the perceived intensity of the stimuli, then the 200 Hz condition should still show significantly better spatial acuity than the 50 Hz condition (hypothesis 3). We found that the spatial acuity for 50 Hz (32.91 ± 7.57 mm) and 200 Hz stimuli (33.35 ± 12.68 mm) was not statistically different when matched for perceived intensity (t_11_ = −0.03; *p* = .979; d_z_ = −.008). Given that the experiment was designed to allow asserting the null hypothesis (i.e., no difference in spatial acuity for 50 Hz stimuli and perceptually matched 200 Hz stimuli), the non-significant result was further investigated through a Bayesian t-tests. The analysis showed that the data were more likely under hypothesis 1 (B_01_ = 3.48, error % = 0.021) than under hypothesis 3. Crucially, hypothesis 2, proposing a higher acuity for the 50 Hz stimuli based on the classical physiological findings on the higher RA innervation density, was still not supported by the data, as the data were also more likely under hypothesis 1 than hypothesis 2 (B_01_ = 3.41, error % = 0.011). This suggests that even after matching the perceived intensity of the stimuli, the 200 Hz stimuli showed a spatial acuity virtually identical to that of the 50 Hz stimuli.

**Figure 2.**
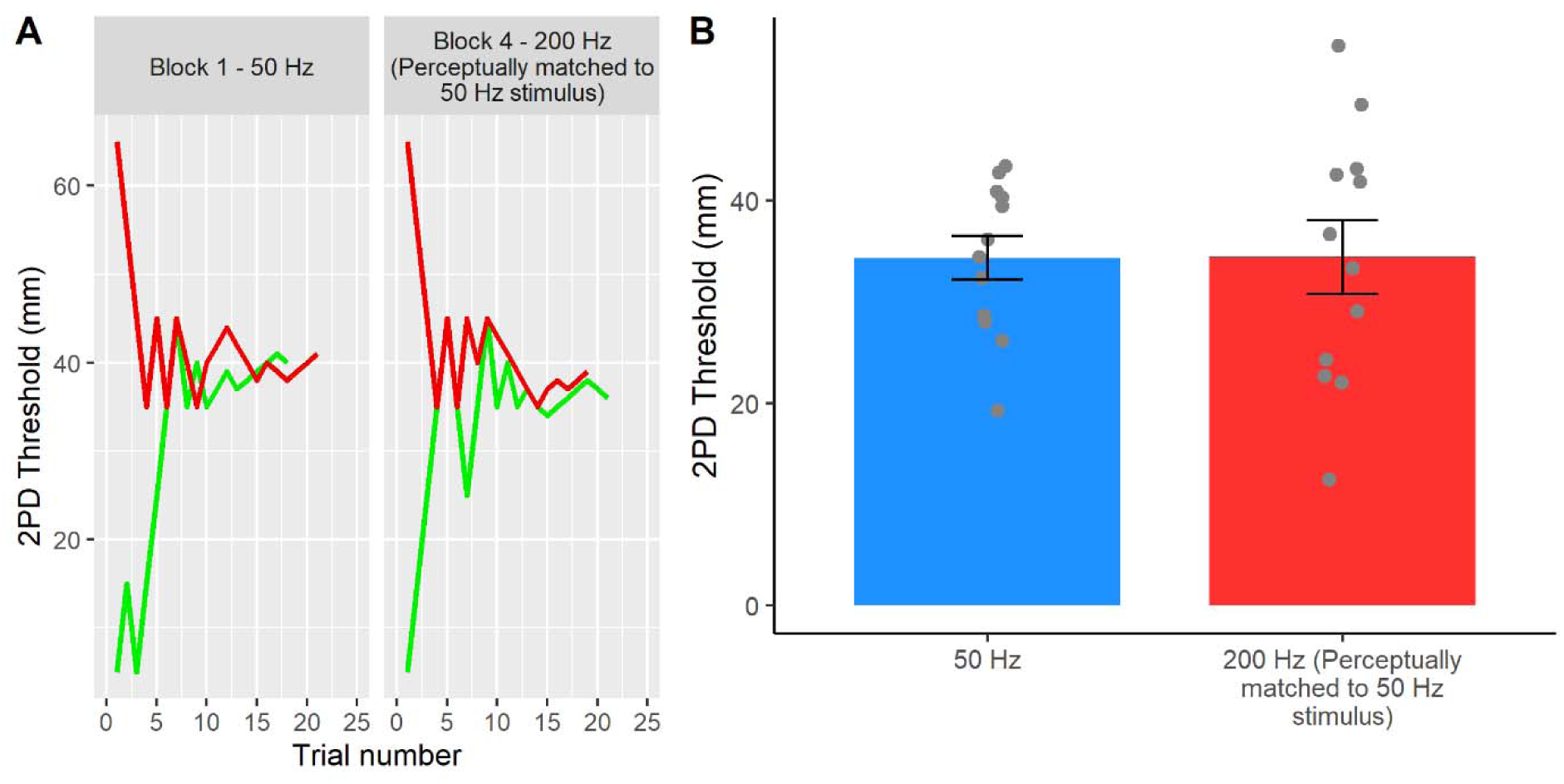
Results from Experiment 2. **A. Staircase data from the 50 Hz and perceptually matched 200 Hz conditions in a representative participant (P# 12, blocks 1 and 4).** In this experiment on the palms, the 50 Hz and 200 Hz stimuli were set to have the same perceptual detectability before 2PDT measurement. Red and green lines show participants responses to descending and ascending staircases, respectively. **B. Two-point discrimination thresholds.** Balancing for subjective intensity showed that, contrary to previous reports, 50 Hz flutter and 200 Hz vibration are perceived with similar spatial resolution. This result was supported by Bayesian analyses (see text). Error bars represent the SEM.

### Does the detectability of the stimuli mediate the effect of perceived intensity and frequency on spatial acuity?

Given that the results yielded by Experiments 1 and 2 were inconsistent with our original hypothesis and the classical Literature, we designed and preregistered Experiment 3 as a replication study (see preregistration at https://osf.io/vf7jy), to investigate the effects from Experiments 1 and 2 in a fully within-participants design. The experiment had three main conditions of ultrasound stimulation: 50 Hz, physically matched 200 Hz, and perceptually matched 200 Hz. Importantly, besides the intensity matching task (Experiment 2) and the 2PD task (Experiments 1 and 2), Experiment 3 also featured a signal detection task aimed to measure the detectability of each of the three ultrasound stimuli. Participants received an equal number of trials where an ultrasound stimulus was either present or absent and were asked to report if they felt a stimulus or not. This task thus provided a measure of participants’ sensitivity (d’) and response bias (C) for each of the three tested conditions.

A series of preregistered planned comparisons between the three conditions (see Statistical Analyses and the preregistration at https://osf.io/vf7jy) showed that Experiment 3 successfully replicated the results of Experiments 1 and 2. In particular, the participants’ 2PD threshold was significantly lower for the 200 Hz stimuli (24.30 ± 10.16 mm), indicating better performance, compared to the 50 Hz stimuli (27.78 ± 7.55 mm) when the two stimuli were physically-matched in terms of intensity (t_19_ = 1.89; *p* = .039; d_z_ = .418), thus replicating the findings of Experiment 1. Moreover, the perceived intensity of the 200 Hz stimulus was again significantly higher than that of the 50 Hz stimulus (48.95 ± 7.58 %; t_20_ = −30.10; *p* < .001; d_z_ = −6.731), as in Experiment 2. Most importantly, when the 50 Hz and 200 Hz stimuli were now perceptually matched for intensity, the 2PD thresholds became virtually identical in the two conditions (perceptually matched condition: 29.94 ± 10.76; t_19_ = −1.10; *p* = .143; d_z_ = −.246; B_01_ = 2.53, error % = 0.016), thus replicating the findings from Experiment 2. The within-participants design of Experiment 3 also allowed to directly compare the 2PD thresholds in the physically matched and the perceptually matched 200 Hz conditions. This allowed us to test whether changes in the physical intensity of a frequency-specific stimulus have an effect on the participants’ spatial acuity. A direct planned comparison confirmed this hypothesis, showing a significant difference (t_19_ = −2.41; *p* = .013; d_z_ = −.540). That is, when the intensity of the 200 Hz stimulus was reduced, in our case by around 50 %, so as to match the 50 Hz stimulus, the participants’ 2PD threshold was significantly higher (i.e., worse acuity). This suggests that spatial acuity depends on stimulus intensity.

Finally, to investigate whether the effect of frequency and perceived intensity on spatial acuity was mediated by the detectability of the stimuli, we ran a one-way repeated measures ANOVA on the tactile detection data for each of the three conditions (50 Hz, physically matched 200 Hz, and perceptually matched 200 Hz). The analysis of the participants’ response bias (C) was non-significant (*F_2,38_* = 2.40; *p* = .104; _η_ *_p_* = 112). Instead, the analysis on the participants’ sensitivity (d’) showed a significant main effect of ultrasound condition (*F_1.29,24.50_* = 15.73; *p* < .001; _η_ *_p_* = 453). A series of Bonferroni-corrected pairwise comparisons showed that the sensitivity to the 50 Hz stimulus (2.40 ± 1.42 a.u.) was significantly lower than both the physically (3.53 ± 0.73 a.u.; t_19_ = −4.52; p < 0.001; dz = −1.01) and the perceptually matched 200 Hz stimulus (3.28 ± 0.79 a.u.; t_19_ = −3.59; *p* = .002; d_z_ = −.803), suggesting that spatial acuity for 50 Hz stimuli might be low due to poor detectability of those stimuli. Crucially, however, the detectability of the perceptually matched 200 Hz stimulus was not statistically different from the physically matched 200 Hz stimulus, even though the 2PD thresholds for those conditions was significantly different. This result suggests that the relationship between physical/perceived intensity and spatial acuity is not simply explained by higher detectability. In fact, even when the stimuli are equally detectable, the perceived intensity of the stimuli still influences spatial acuity.

**Figure 3.**
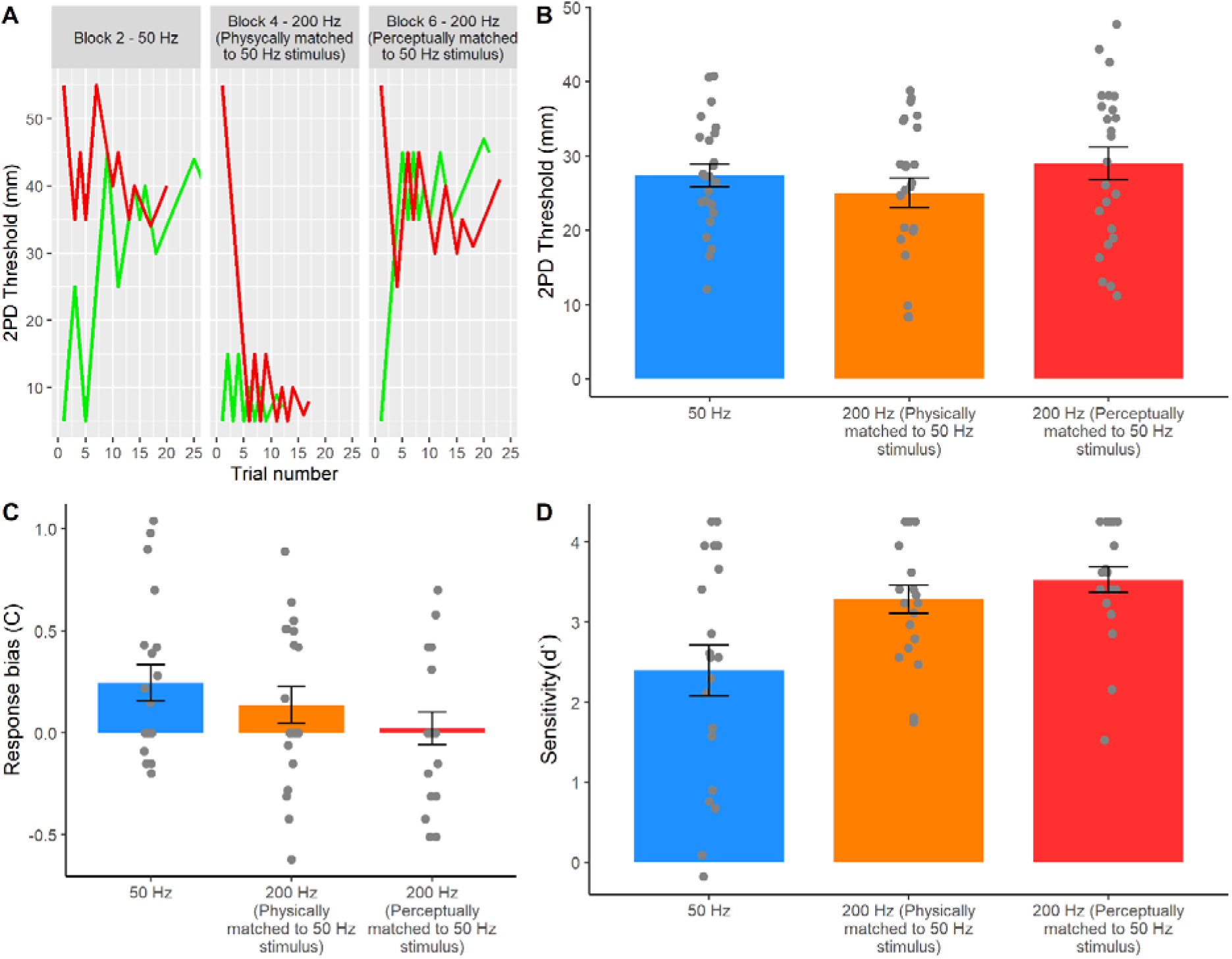
Results from Experiment 3. **A. Staircase data from the 50 Hz condition and the physically and perceptually matched 200 Hz conditions in a representative participant (P# 26, blocks 2, 4, and 6).** This preregistered replication experiment (https://osf.io/vf7jy) tested the participants’ 2PD thresholds for 50 Hz and physically and perceptually matched 200 Hz stimuli. Red and green lines show responses to descending and ascending staircases, respectively. **B. Two-point discrimination thresholds.** The 2PD threshold results from Experiments 1 and 2 were successfully replicated. The 200 Hz stimulus produced lower 2PD thresholds (i.e., better spatial resolution) than the 50 Hz stimulus when the two conditions had equal physical intensity. However, when the two conditions were matched to have equal perceived intensity, 50 Hz and 200 Hz stimuli were perceived with similar spatial resolution. **C. Response bias data.** The second part of Experiment 3 investigated whether the effect of frequency and perceived intensity on spatial acuity was mediated by the detectability of the stimuli. Differences between conditions were not significant. **D. Sensitivity data.** Sensitivity to the 50 Hz stimulus was significantly lower than to both the physically and the perceptually matched 200 Hz stimulus. Error bars represent the SEM.

### Do frequency-specific differences in 2PDT reflect differences in perception, or differences in skin mechanical response to stimulation?

Somatosensory research typically controls the stimulus delivered to the skin surface but often ignores the mechanical propagation through the skin to the sensory receptor (but see^24,25,34,35^ for exceptions). The *effective* stimulation to the receptor is generally unknown. One may therefore ask whether the 2PDTs measured for 200 Hz and 50 Hz stimuli really reflect the perceptual acuity of the corresponding submodality channels, or merely the way the ultrasound stimulus mechanically interacts with the skin, leading to frequency-specific differences in effective stimulation.

To test whether the results from Experiments 1-3 represent genuine perceptual processes rather than artefacts of varying mechanical response properties of skin, Control Experiment 4 used a laser doppler vibrometer (LDV) to measure the precise peak-to-peak skin displacement and propagation waves induced by each of the stimuli we tested. First, we tested if the average peak-to-peak skin displacement differed between 50 Hz and 200 Hz ultrasound stimulations. As a check, we confirmed that 50 Hz stimuli was perceived as weaker (53.10 ± 6.93 %) than 200 Hz stimuli of the same physical amplitude, thus replicating our results from Experiments 2 and 3. We reasoned that this difference in perceived intensity might reflect the mechanical response of the skin to different ultrasound frequencies, leading to different *effective* stimulation of the RA and PC receptors. In particular, although we fixed the acoustic pressure (i.e., physical intensity) of our ultrasound stimuli to be the same for our 50 Hz and 200 Hz stimuli, the effective stimulation at the receptor might still be greater for 200 Hz than for 50 Hz stimuli because the skin might resonate more to 200 Hz that to 50 Hz stimuli, resulting in better effective stimulation for 200 Hz than 50 Hz. In that case, our findings of higher perceived intensity for 200 Hz stimuli physically matched to 50 Hz stimuli would simply reflect a peripheral, non-neural effect due to skin mechanics. We therefore used LDV to measure the actual peak-to-peak displacement of the skin caused by 50 Hz ultrasound stimuli and by physically-matched and perceptually-matched 200 Hz stimuli.

Contrary to the differential mechanical resonance hypothesis, we found that in all the four participants tested, the average peak-to-peak skin displacement was in fact greater in the 50 Hz condition (mean ± SD: 7.61 ± 1.87 µm) than in the physically-matched 200 Hz condition (4.13 ± 0.85 µm). Thus, although across Experiments 2-4 the perceived intensity of the 50 Hz stimulus was much lower than the perceived intensity of a 200 Hz stimulus having the same physical amplitude, the actual skin displacement induced by the 50 Hz condition was on average 184% of the displacement in the 200 Hz condition.

Next, we quantified the average skin displacement due to a 200 Hz stimulus that was perceptually matched in intensity to the 50 Hz stimulus and found this to be 0.92 ± 0.36 µm – just 12% of the displacement in the 50 Hz condition. These results clearly rule out the possibility that the low perceived intensity of 50 Hz stimuli simply reflects poor resonance of the skin to 50 Hz stimulation. Rather, they suggest a frequency-dependent non-linear relationship between physical skin displacement and perceived intensity.

**Figure 4.**
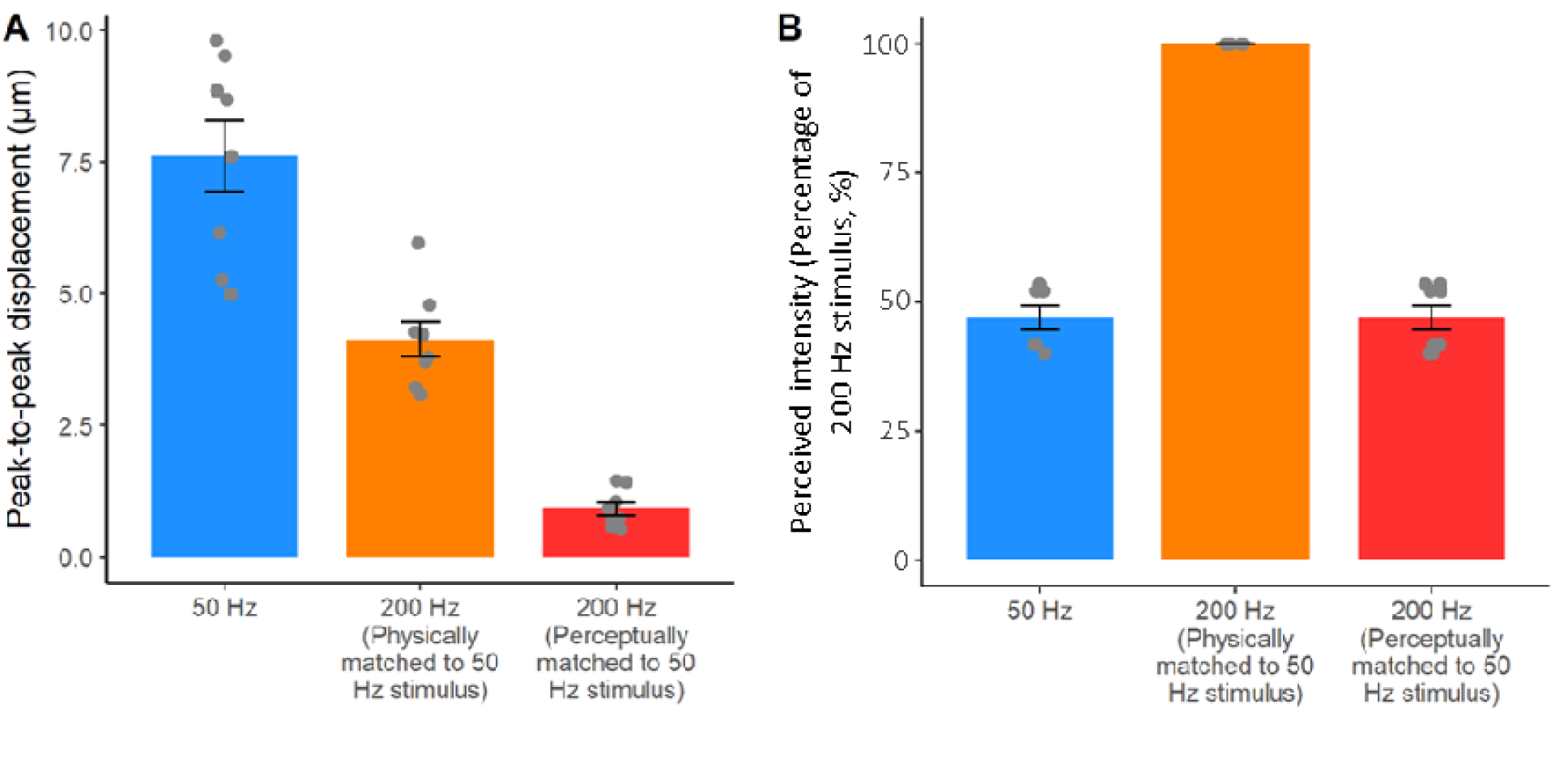
Results from Experiment 4. **A. Peak-to-peak skin displacement caused by the three ultrasound stimuli.** Contrary to a differential mechanical resonance hypothesis, we found that the average peak-to-peak skin displacement was greater in the 50 Hz condition, compared to the 200 Hz conditions. **B. Perceived intensity for the same stimuli in A.** The 50 Hz stimuli produced the largest skin displacement, but were perceived as having less than half the intensity of 200 Hz stimuli that were matched for physical intensity, and that produced much smaller skin displacement.

Next, we tested whether the effects of frequency and intensity on spatial acuity might be due to the interference created by the two mechanical propagation waves on the skin that inevitably arise when two points are stimulated simultaneously^24,25^. The 2D and 1D plots of the skin displacement induced by the two points at the participants’ 2PD threshold show that the displacement of the skin in the region between the two stimulated points was minimal. Moreover, the skin displacement was on average lower when the two points were at 2PD threshold (3.93 ± 2.97 µm), compared to a suprathreshold condition (4.51 ± 3.20 µm) (Figure 5 and Movie S1), thus refuting the idea that the observed 2PD thresholds could be influenced by skin mechanics phenomena such as surface wave propagations. These data also confirm the success of our attempts to design ultrasound stimuli with minimal side-lobes for the specific purpose of acuity testing (see Methods).

**Figure 5.**
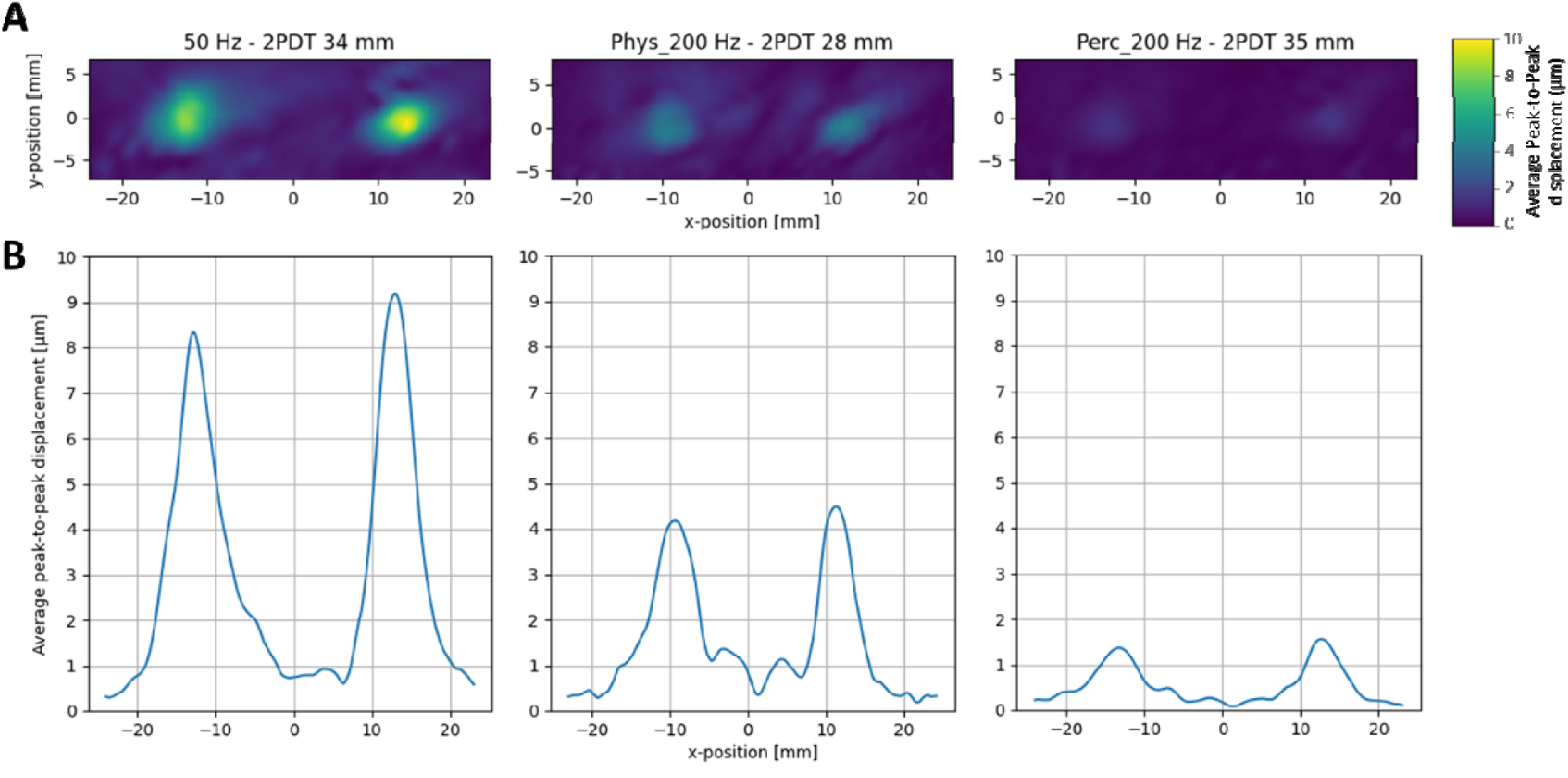
A. Heatmaps of the peak-to-peak skin displacement (μm) in a representative participant. LDV recording were obtained in each stimulation condition at its 2PD threshold. **B. Cross-section of the two-point maxima.** The plots show the 1D cross-section of the same data shown in A.

## Discussion

Spatial acuity is a fundamental property of the somatosensory system, but it has been studied using exclusively mechanical pressure stimuli. This means that spatial acuity has been properly characterised only for one of the many mechanoreceptive channels constituting the somatosensory system (i.e., SAI). We used a novel, contactless ultrasound stimulation to systematically investigate the spatial acuity for the frequency-resolved mechanoreceptive channels (RA, PC). Classically, the PC system is thought to transmit little or no spatial information, due to its extreme sensitivity, low innervation density, deep receptors, and large, indistinct receptive fields.

Contrary to this hypothesis, across four experiments, we found that spatial acuity for 200 Hz stimuli (preferentially activating PC afferents) was at least as good as the acuity for 50 Hz stimuli (preferentially activating RA afferents). Moreover, in Experiments 1 and 3 spatial acuity was better for 200 Hz stimuli than for 50 Hz stimuli of equal stimulus energy. We showed that this surprising result was partly due to the fact that 50 Hz stimuli feel less intense than 200 Hz stimuli of equal stimulus energy. Therefore, it is possible that 50 Hz acuity judgement may be poor because stimuli are faint. Therefore, Experiment 2 equalised the perceived intensity of the stimuli, and then found that tactile acuity for 50 Hz and 200 Hz stimuli was essentially equivalent. More generally, sensitivity and detectability are important factors to take into account in quantifying spatial resolution.

The low sensitivity of the 50 Hz receptor might reflect poor mechanical transmission of 50 Hz stimuli by the skin to the receptor or might reflect a truly low sensitivity of the Meissner receptor itself, relative to the Pacinian. We used LDV measurements to address this question, and conclusively ruled out the first possibility. We showed that 50 Hz stimuli produce larger peak-to-peak skin displacement than 200 Hz stimuli of the same physical intensity, and much more skin displacement than 200 Hz stimuli of the same perceived intensity. We also showed that the spatial spread of mechanical energy across the skin is comparable between 50 Hz and 200 Hz stimuli, with no evidence for differential smearing at intermediate locations between two-point stimuli leading to confounds in acuity measurement. For these reasons, we can conclude that differences in sensitivity and acuity between frequency-specific submodalities reflect genuine features of the corresponding neural channel, and not merely differences in effective stimulation or coupling between the stimulator and the receptor. Thus, our data both confirms the previously reported high sensitivity of the PC channel relative to the Meissner/RA channel, and now adds the novel finding that the PC channel has a higher spatial acuity than previously thought.

Classically, the PC channel of human mechanoreception was identified by its high sensitivity to high-frequency sinusoidal vibrations. More recently, Birznieks et al.^36^ used low amplitude pulsatile stimuli to activate the PC channel. Because the amplitude of their stimuli was well below the RA/Meissner channel threshold, they assumed that any resulting percept was due to the PC channel. They showed that a perceptual experience could be evoked from the PC channel for pulses at frequencies down to 6 Hz. Moreover, these stimuli produced an experience of vibration that was continuous with the experiences of vibrations targeting the RA/Meissner channel, rather than qualitatively different from it. These results question the classical view of different receptor types being associated with distinct sensory qualities and distinct central projections, and instead suggest a common perceptual dimension of frequency that is encoded by the pattern of activation across multiple afferents. Our study used stimuli designed to target a single receptor type and did not explore the frequency continuum. In addition, we have focussed on the spatial percepts generated by frequency-targeted stimuli, in contrast to the more extensive literature on temporal aspects of perception. However, both sets of results indicate that the role of the PC channel in human somatosensory perception may be wider than previously thought, given that PCs encode a surprising amount of temporal information^36^ and also spatial information (present results).

Our results have special implications for haptic applications. Mid-air haptics using ultrasound stimuli to produce tactile experiences without mechanical contact between skin and stimulator. These devices cannot provide steady pressure or low-frequency stimulation, and therefore cannot currently produce experiences such as hand-object interactions, grasping etc. For this reason, recent interest has often focussed on temporal pattern perception, including texture perception^37^. Our results suggest that ultrasound stimulation can nevertheless elicit spatial percepts, and that transmission of spatial patterns is a useful application goal for mid-air haptics.

Our study has a number of limitations. Some of these result from the ultrasonic stimulation mechanism and can be considered as corollary disadvantages that accompany ultrasound’s unique capability to provide frequency-resolved skin stimulation. For example, ultrasound stimuli are weak, and the contact forces and skin displacement in our studies are therefore lower than those conventionally used in spatial resolution studies. Some participants were excluded because of difficulty in detecting some stimuli, particularly at lower frequencies.

Other ultrasound tactile studies report similar exclusions^38^. As a consequence of these exclusions, our results may not generalize well to the entire population. More generally, the relation between stimulus intensity and spatial acuity is not well understood. Our own data show a relation between sensory detection, which depends mainly on stimulus intensity, and spatial acuity. We therefore speculate that stronger stimulation might reveal even better spatial acuity of the PC channel. On the other hand, several studies suggest that spatial information and force/pressure information are processed in separate brain pathways^39,40^.

Second, ultrasonic stimulation provides a focal point of stimulation that is much larger than the probes conventionally used in acuity testing. Our LVD recordings indicate that each point of stimulation is ~6 mm in diameter (see Methods). The relatively low spatial resolution of stimulation inevitably leads to relatively high estimates of two-point threshold, due to the spread of stimulation from the centre or each focal point. However, focal point size is roughly constant across stimulation frequencies, as our LDV measures show. Therefore, this limitation does not undermine our finding of surprisingly good spatial resolution of the PC channel. A useful future research exercise could involve directly measuring the relation between stimulus spread and two-point threshold spatial acuity estimates for a range of different stimulation devices in the same participants.

To conclude, we have taken advantage of the frequency-resolved spatially-controlled stimulation provided by ultrasound arrays to measure the spatial resolution of human tactile perception at the distinct 50 Hz and 200 Hz frequencies associated with the RA/Meissner and PC channels respectively. We found that spatial acuity for 200 Hz stimuli is comparable to that for 50 Hz stimuli matched for subjective perceptual intensity. This finding is interesting, given classical anatomical and physiological reports of low innervation density, and large, indistinct receptive fields of PCs. Experience in mid-air haptics teaches that 200 Hz stimuli are among the most readily detected by human participants in interaction tasks. Our result therefore favours the use of spatially-patterned stimuli, as well as temporally-patterned stimuli in applications using mid-air haptics for human interactions.

**Figure 6.**
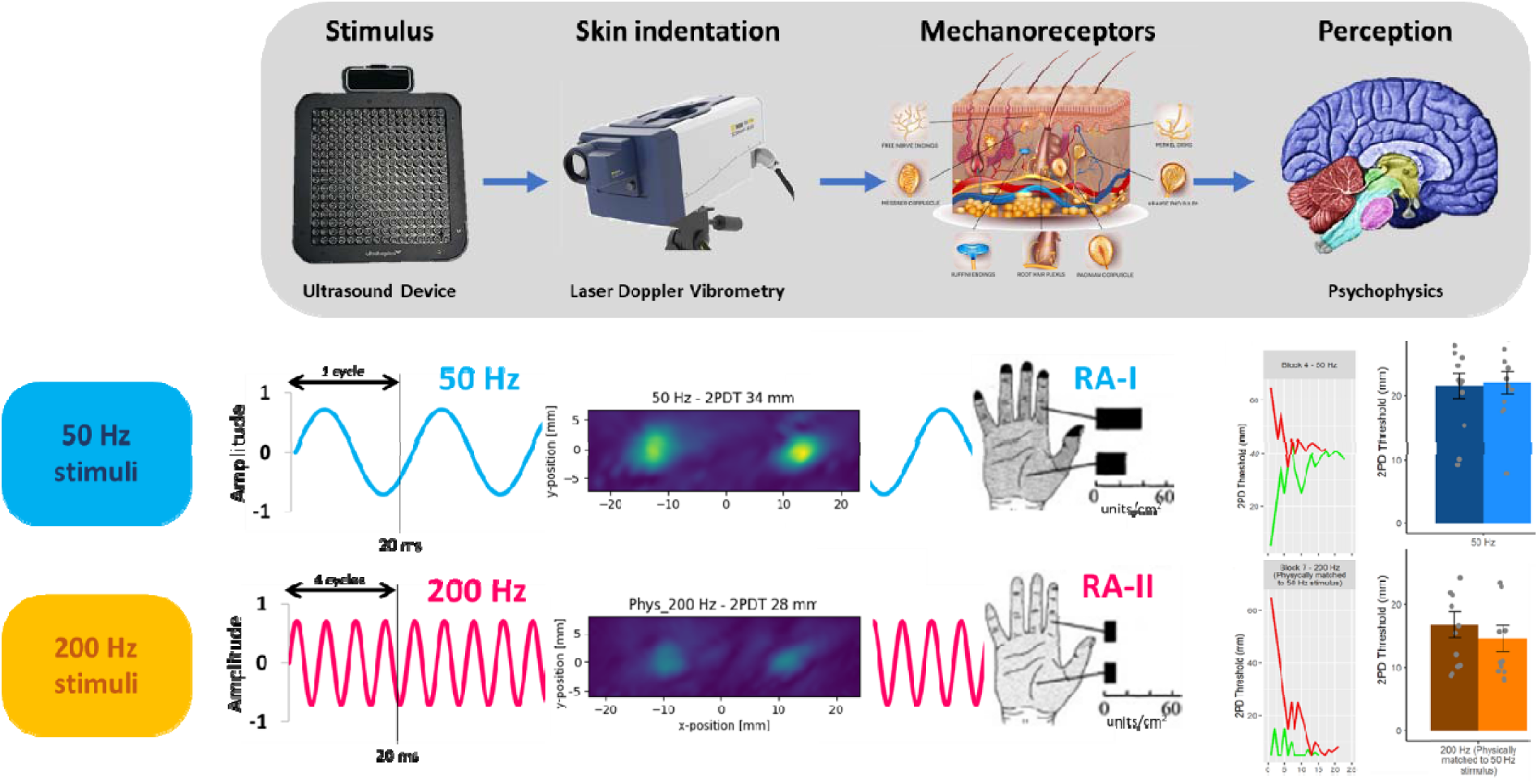
Schematic showing the logic of inter-channel comparisons for perceptual studies. The stimulator device can be programmed to deliver a desired stimulus to the 50 Hz or the 200 Hz channel. Differences betwee the percepts yielded by the two channels could reflect mechanical differences in effective stimulation (for example due to differences in the skin indentations caused by the two stimuli), or neural differences in transduction/transmission by the corresponding mechanoreceptors/afferents. LVD measures of skin indentation confirm that the inter-channel differences that we observed in perception were unlikely to be due to differences in effective stimulation and were therefore presumably neural in origin.

## Acknowledgments

This study was supported by the European Union’s Horizon 2020 research and innovation programme under grant agreement No 101017746.

## Authors’ contributions

Conceptualization: A.C. and P.H.; methodology: A.C., W.F., and P.H.; software, validation, formal analysis, and visualization: A.C. and W.F.; resources: P.H.; data curation: A.C. and W.F.; writing – original draft: A.C.; writing – review & editing: A.C., W.F., and P.H.; supervision, project administration, and funding acquisition: P.H.

## STAR Methods

### Resource availability

#### Lead contact

Further information and requests for resources should be directed to and will be fulfilled by the lead contact, Antonio Cataldo.

#### Material availability

This study did not generate new unique material.

#### Data and Code availability

All data needed to evaluate the conclusions in the paper are present in the paper and/or the Supplemental Information.

This paper does not report original code.

Any additional information is available from the lead contact upon request.

### Experimental model and subject details

#### Participants

The sample size for Experiments 1-2 (n = 12) was decided a-priori on the basis of a power analysis on the results of a previous similar experiment^21^ (see preregistration at https://osf.io/jkd8h). In that study, Tannan and colleagues tested the 2PD thresholds of four participants for static stimuli, low-frequency (25 Hz), and high-frequency vibrotactile stimuli (200 Hz). Although the authors did not report the effect size for the main effect of frequency, from the means and SD of each condition reported in their Figure 7, we estimated that the effect size for the difference between low- and high-frequency conditions was Cohen’s dz = 3.39, considered to be extremely large using Cohen’s criteria ^41^. With an alpha = 0.05 and power = 0.95, the projected sample size indicated to demonstrate differences in 2PD thresholds based on different vibration frequencies was 4 participants^42^. Nonetheless, we decided to set a sample size of n = 12 to account for the differences between our setup and the one used by Tannan and colleagues^21^ (i.e., mechanical vs. ultrasound stimulation).

The sample size for Experiment 3 was also preregistered (see preregistration at https://osf.io/vf7jy) and estimated a-priori through a power analysis on the results Experiment 1. In Experiment 1 the effect size for the difference in 2PD thresholds between a 50 Hz stimulus and a physically-matched 200 Hz stimulus was Cohen’s dz = .846, considered to be large using Cohen’s criteria^41^. With an alpha = 0.05 and power = 0.80, the projected sample size indicated to demonstrate differences in 2PD thresholds based on different vibration frequencies was 14 participants. Nevertheless, we set a stopping rule of 18 participants to counterbalance the order of the three conditions (i.e., six permutations) across participants. Two extra participants were tested and included in the analyses because they had already been recruited prior to the reaching of the stopping rule. Therefore, the final sample size for Experiment 3 was n = 20. Finally, the sample size for Control Experiment 4 (n = 4) was based on an opportunity sample. The low n was due to the technical challenges involved in recording LDV data, including long (~3 h+) experimental sessions and prolonged immobilisation of the left hand and arm.

A total of 57 right-handed healthy participants (35 females; mean age ± SD: 25.0 ± 5.0) were originally recruited. Based on preregistered exclusion criteria (see preregistration at https://osf.io/jkd8h, Figure S5, and Dataset S1), three participants were excluded because they produced a false alarm in more than 40 % of the catch trials (i.e., they reported a two-point stimulus when only one point was delivered). Three more participants were excluded because they could not feel a clear sensation in at least one of the stimulation conditions. Finally, two more participants were excluded because they did not follow the instructions and one could not be tested due to technical issues. The experimental protocol was approved by the Research Ethics Committee of University College London and adhered to the ethical standards of the Declaration of Helsinki. All participants were naïve regarding the hypotheses underlying the experiment and provided their written informed consent before the beginning of the testing, after receiving written and verbal explanations of the purpose of the study. All participants received a monetary compensation (£8 per hour) for their involvement in the study. The hypotheses, procedures, and analyses of Experiments 1 (https://osf.io/jkd8h) and Experiment 3-4 were pre-registered (https://osf.io/vf7jy).

### Method Details

#### General Setup

Figure S1 shows the experimental setup for Experiments 1-3. Participants sat at a desk, resting their left elbow on an articulated armrest support (YANGHX, model 4328350928, China) and their left hand palm down on a plastic support, in correspondence of a 10 x 10 cm aperture. Participants’ view of their left hand was blocked by a fixed vertical plywood screen throughout the entire experiment. Airborne ultrasound stimulation was delivered through a STRATOS^TM^ Explore device (Ultraleap, UK), located 12 cm below the participant’s hand. We identified a technical error which placed the focal point for the single-point “catch” trials in Experiments 1 and 2 at 10 cm rather than 12 cm from the participants’ hand. This resulted in the catch trials having a lower intensity compared to the other trials. These catch trials are not included in the calculation of 2PD, and only used to screen for participants who habitually respond “two” to every stimulation – implying infinite spatial resolution^12^ (see also our preregistration at https://osf.io/jkd8h). Therefore, the unintended low intensity of these catch trials in Experiments 1 and 2 should not affect our 2PD estimates, though it might have affected exclusion of some participants. The STRATOS^TM^ Explore is a computer-controlled device composed by an array of 16 x 16 transducers which employs focussed ultrasound (40 KHz) to project discrete tactile points directly onto the participants’ skin^28^. The amplitude of the focal point generated by the device (~6 mm diameter) can be spatiotemporally modulated to produce vibrotactile patterns of different shapes and frequencies^30^. This setup allowed us to deliver single vs two-points contactless, frequency-resolved vibrotactile stimuli.

To measure the peak-to-peak displacement produced by the ultrasound stimulation on the participants hands, in Experiment 4 we used a LDV PSV-500-Scanning-Vibrometer from Polytec. The LDV head was aimed at the participant hand at an angle of 45 degrees. This angle was later accounted for in the data analysis by d = d′/sin(θ), where d is the estimated perpendicular displacement and d′ is the measured displacement and θ is the acute angle between the laser and the horizontal surface. The participants laid flat on a dentist chair, their arm resting on an optical table (Thor labs) and supported by a vacuum cushion. The subject palm was facing upwards and the ultrasound device (Ultrahaptics STRATOS^TM^ Explore) downwards (see Figure S2).

All the software for the experimental tasks were coded in Python 3.6^43^.

#### Creation and validation of two-points stimuli

To produce the different mid-air haptic stimuli, i.e., pair of points, we used a STRATOS Explore from Ultraleap that consisted of 256 ultrasonic transducers operating at 40kHz. The device API implements a focusing algorithm based on eigen vector^30^ and allows users to simply create one or more mid-air haptic focal points at the desired positions and with the desired amplitude (see Figure S5). We defined the size of a focal point using the Full Width Half Max (FWHM) metric. This metric computes the maximum distance across the focal point where the acoustic pressure is greater or equal to half of the peak acoustic pressure. In our experiments, the focal point was set to an FWHM of 6 mm diameter. Thus, we defined two focal points defined whose amplitude fluctuated according to a sinusoidal signal at a rate of either 50Hz or 200Hz, depending on the condition. To make the most of the device output, the focal points amplitude signals were defined with opposite phased. In other words, when a focal point amplitude was maximal the other point amplitude was minimal. However, the symmetry within the rectilinear arrangement of the transducers can yield to the creation of grating lobes^44^. To avoid such grating lobes occurring in between the pair of focal points, we defined a third focal points which amplitude was set to 0 throughout the stimuli duration. In summary, the amplitude of the focal points was defined as:

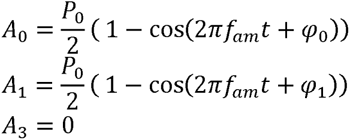

Where *P*_0_ is equal to 2000Pa, *φ*_0_ and *φ*_1_ are equal to 0 and π, respectively, and *f_am_* is equal to 50Hz or 200Hz depending on the condition.

The focal points were targeted onto the palm which was placed 120mm directly under the haptic device such that the applied force was normal to the surface of the hand. The points coordinates were set to (−s/2; 0; 120) mm, (s/2; 0; 120) mm, and (0,0,120) mm, where s is the spacing between the 2-points.

#### Experimental design and procedure

The main dependent variable across all experiments was the participants’ 2PD threshold, defined as the minimal distance required to detect two simultaneous adjacent ultrasound points with a 50% accuracy.

#### Experiment 1: is spatial acuity affected by the frequency of the stimuli?

Experiment 1 aimed to test whether the frequency of a vibrotactile stimulus affected the participants’ 2PD threshold (see preregistration at https://osf.io/jkd8h). The experiment had a 2 (Frequency: 50, 200 Hz) x 2 (Skin region: palm, index finger) within-participants factorial design. In each of the four factorial combinations, the participants’ 2PD threshold was tested twice, for a total of eight blocks. The presentation order of the blocks was counterbalanced and pseudo-randomised across participants such that the levels of the Skin region factor were always interleaved. This manipulation was intended to minimise as much as possible the sensitisation/fatigue of the mechanoreceptors (see for example^12^). Participants took short breaks between blocks and were asked to move and stretch their body so to prevent any discomfort due to prolonged testing. Each of the eight blocks lasted about five minutes, for a testing session of approximately 75 minutes.

In each block, the participants performed a 2PD adaptive staircase procedure for simultaneous stimuli based on the methods described by Mancini and colleagues^12^. Each block started with a calibration phase where a two-points stimulus at the largest distance (65 mm) was delivered on the tested skin region (palm or index finger). The participants were asked to adjust the position of their hand/index finger to make sure that both points were centred and clearly perceivable. Then, a single (~20% of trials) or a two-points (~80% of trials) ultrasound stimulus was delivered for 3 s. The location of the single point stimulus on the tested body part was randomised across three different positions (distal, middle, and proximal) to ensure that the position of the stimulation was not indicative of whether the trial contained a single- or two-pointed stimulus. Two consecutive beeps signalled the beginning and the end of the stimulation. After the second beep, the participants reported whether they felt one or two points by pressing one of two keys with their right hand. The next trial started after an intertrial interval of 500 ms from the participant’s keypress

The distance between the two simultaneous points was adaptively adjusted according to the participants’ response to the previous trial. A correct identification produced a decrease in the two-points distance, while an error produced an increase in the two-points distance. Two randomly-interleaved staircases were used in each block, one starting from the smallest testable distance (5 mm) and the other starting from the largest testable distance (65 mm), in order to minimise the participant’s expectations about incoming stimuli^45^. Increasingly smaller step-sizes (10, 5, 2, and 1 mm) were used after the first two reversals, to provide a fast convergence toward the psychophysical thresholds^45^. The staircase procedure ended after reaching ten reversals. The average distance of the last eight reversals was taken as the 2PD threshold for that specific block. The final estimate of each participant’s 2PD threshold was defined as the average of each repetition block testing the same condition in a given experiment (see Experimental design).

#### Experiment 2: is the effect of frequency on spatial acuity meditated by the perceived intensity of the stimuli?

Experiment 2 aimed to test whether the two vibrotactile frequency patterns used in Experiment 1 had difference perceived intensity, and whether this difference mediated the results of Experiment 1. The experiment employed a procedure similar to Experiment 1, but it also featured an intensity matching task prior to the 2PD testing. This task provided the point of subjective equality (PSE) or perceived iso-intensity levels for the two tested frequencies (50, 200 Hz). Given that the Skin region factor in Experiment 1 was not significant, we dropped that factor in Experiment 2 to allow for a shorter testing session. Thus, Experiment 2 had a single factor with two levels (Frequency: 50, 200 Hz). The participants’ PSE for the intensity of the two vibrotactile stimuli was tested only once (~7 min), while the 2PD thresholds were tested three times in each condition, for a total of six blocks, each lasting ~ 5 min. Short breaks were provided between blocks.

The intensity matching task was based on a 2-AFC adaptive staircase testing participants’ intensity perception rather than spatial acuity. In each trial, participants received two consecutive ultrasound stimulations. Each stimulation was made of two-15 mm-points and lasted for 1 s. The distance between the two points was fixed to 15 mm based on the results of Experiment 1, where the smallest 2PD threshold across conditions was in average 14.54 ± 6.96 mm. Each stimulus was associated with a beep. After the second beep, participants were asked to report which beep contained the strongest vibrotactile stimulus. The first stimulus in the trial (i.e., reference stimulus) was a 50 Hz stimulus at the highest possible intensity allowed by the STRATOS^TM^ Explore device. The second stimulus (i.e., comparison stimulus) was a 200 Hz stimulus whose intensity varied depending on the participants’ response. Randomly interleaved ascending and descending staircases were also used in this task. Increasingly smaller step sizes (20, 10, 5% of strongest intensity) were used. Each staircase ended after ten reversals. The average of the last eight reversals was used as an estimate of the perceived intensity ratio between the 50 Hz and the 200 Hz stimulus.

#### Experiment 3: is the effect of perceived intensity on spatial acuity meditated by the detectability of the stimuli?

Experiment 3 aimed to replicate and expand the findings from Experiment 1 and 2 in a full within-participants design. The experiment had three main conditions of ultrasound stimulation: 50 Hz, physically matched 200 Hz, and perceptually matched 200 Hz. These conditions were selected to answer four specific research questions (see preregistration at https://osf.io/vf7jy):

1. Is the 2PD threshold of a vibrotactile stimulus affected by its frequency?
2. Is the perceived intensity of the vibrotactile stimulus affected by its frequency?
3. Is the effect of frequency on spatial acuity mediated by the perceived intensity of the stimuli?
4. Is the effect of perceived intensity on spatial acuity mediated by the detectability of the stimuli?

Participants performed an intensity matching task at the beginning of the experiment to establish the intensity of the 200 Hz stimulus perceptually matched to the 50 Hz stimulus. Instead of using a two-point stimulus for the intensity matching task as in Experiment 2, here we used a single central point. The reason for this choice was to avoid any spatial summation in this purely intensive task. A small pilot study (n = 8) conducted prior to the beginning of the experiment confirmed that delivering a single or a double point as a stimulus for the intensity-matching task did not produce statistically different results (see Figure S3). The intensity matching task was followed by a 2PD task and then by a signal detection task. The 2PD task was identical to Experiment 1 and 2, except that each stimulus lasted 1 s and that the largest two-point distance tested was 55 ms rather than 65 mm. Both these changes were justified by the aim of keeping the duration of the experimental session within ~75 min, despite the higher number of tasks performed.

The signal detection task aimed to test the detectability of each of the three ultrasound stimuli used across Experiments 1 and 2 (50 Hz, physically matched 200 Hz, and perceptually matched 200 Hz). In each of the three blocks, participants were delivered with 30 trials containing a stimulus and 30 empty trials (fully randomised) and were asked to report if the trial contained a stimulus or not. This task thus provided a measure of participants’ sensitivity (d’) and response bias (C) for each of the three conditions.

Participants’ 2PD thresholds were tested twice for each of the three different ultrasound conditions, while their signal detection was tested once for each condition. The presentation order of the three conditions in each task was counterbalanced across participants. Short breaks were allowed between blocks. Each block lasted about five minutes, corresponding to a testing session lasting approximately ~75 minutes including instructions and breaks.

#### Control Experiment 4: is the effect of perceived intensity on spatial acuity due to the propagation of mechanical waves through the skin?

Control Experiment 4 aimed to replicate the methods of Experiment 3 in a small sample of participants while simultaneously characterising the physical peak-to-peak skin displacement induced by the ultrasound stimulus using a Laser Doppler Vibrometry (LDV) procedure. Peak-to-peak skin displacement measures were taken for ultrasound stimuli of 50 Hz, 200 Hz, and perceptually-matched-200 Hz at the participant’s 2PD threshold and at suprathreshold (55 mm) distances.

The LDV used in area scan mode measured the velocity of vibrations across the participants palm with at a sampling rate of 128kHz for a duration of 128ms per scanning locations, corresponding to a sample time of 256 ms. The array was sending a trigger-pulse to the LDV control unit reference channel. The trigger-pulse was sent at the start of a mid-air haptic stimulus, every 128ms (the length of the LDV sample time). The mid-air haptic stimulus was running for 100ms after the trigger time and then stopped. The extra 28 ms without stimulation were there to let the skin come back to a rest state and therefore avoid interference contamination across sample measurements. The displacement was calculated by integrating the recorded velocity time series, prior to integration the velocity data a high pass filter was applied to remove low frequency noise (cut-off of 20 Hz) as well as a low-pass filter to remove frequency not relevant to touch (cut-off 2kHz). Finally, the average peak-to-peak displacement was computed and reported.

This measure allowed us to disentangle whether the difference in spatial acuity and perceived intensity between different frequencies was due to a specific mechanical interference of propagation waves on the skin^24,25^.

### Quantification and statistical analysis

All data except one condition from Experiment 1 (200 Hz-Palm) met the normality assumption (Shapiro-Wilk test: *p* > .054), therefore, parametric tests were used throughout. In Experiment 1, the 2PD thresholds values for each participant were entered in a 2 x 2 repeated measures factorial ANOVA with factors of Frequency and Skin region (see above). Non-significant results were further tested using Bayesian analyses in order to test whether the data were more likely under the null than under the alternative hypothesis^46,47^.

In Experiment 2, we first used a one-sample t-test contrasting the participants’ PSE (representing the perceived iso-intensity of 50 Hz and 200 Hz stimuli) against a test value of 1 (i.e., the physical intensity of the 50 Hz stimulus). This analysis allowed us to investigate whether the perceived intensity of physically iso-intense ultrasound stimuli was modulated by the frequency of the vibrotactile stimulation. Next, we used a paired t-test to compare the participants’ 2PD threshold data in the two frequency conditions (50 Hz vs. perceptually matched 200 Hz stimuli). Given that the experiment was designed to allow asserting the null hypothesis (i.e., no difference in spatial acuity for 50 Hz stimuli and perceptually matched 200 Hz stimuli), the non-significant result was further investigated through Bayesian t-tests.

Experiment 3 was based on four preregistered (https://osf.io/vf7jy) planned comparisons between three tactile conditions (50 Hz, physically matched 200 Hz, and perceptually matched 200 Hz). To replicate the results of Experiment 1 on the effect of vibrotactile frequency on spatial acuity, we ran a one-tailed paired sample t-test on the 2PD thresholds of the 50 Hz stimulus and the physically matched 200 Hz stimulus. Next, to replicate the effect of vibrotactile frequency on perceived intensity (Experiment 2) we ran a one-tailed paired sample t-test on the intensity of the perceptually matched 200 Hz stimulus in the intensity matching task against a test value of 1 (i.e., the intensity of the 50 Hz stimulus). To replicate the effect of perceived intensity on spatial acuity (Experiment 2), we ran a one-tailed paired sample t-test on the 2PD thresholds of the perceptually matched 200 Hz stimulus and the physically matched 200 Hz stimulus, and another one-tailed paired sample t-test on the 2PD thresholds of the 50 Hz condition and the perceptually matched 200 Hz stimulus. Again, given that we predicted a null effect for this latter comparison, we used Bayesian analyses to test whether the data were more likely under the null than under the alternative hypothesis. Finally, to investigate whether the effect of frequency and perceived intensity on spatial acuity was mediated by the detectability of the stimuli, we ran a one-way repeated measures ANOVA on the tactile detection data for each of the three conditions (50 Hz, physically matched 200 Hz, and perceptually matched 200 Hz). Significant results were followed up with appropriate post-hoc analyses. Given that this hypothesis is bidirectional, we again used Bayesian analyses to test whether the data are more likely under the null than under the alternative hypothesis.

Data analyses were ran using R^48^ and IBM SPSS Statistics for Windows v. 25 (IBM Corp., USA). Bayesian analyses were run on JASP v. 0.12.1 (JASP Team 2016, University of Amsterdam).

## Supplemental items legends

**Video S1. (Separate file) Animated gifs of LDV recordings obtained in each stimulation condition at its 2PD threshold, Related to Figure 5**. Video depicting the experimental setup for the hand-driven self-touch condition.

**Data S1. (Separate file) All data necessary for the analyses reported in this study, Related to Figures 1–4.**

