## Supplemental Information for "The Limits of Touch: Spatial acuity for frequency-resolved air-borne ultrasound vibrotactile stimuli"

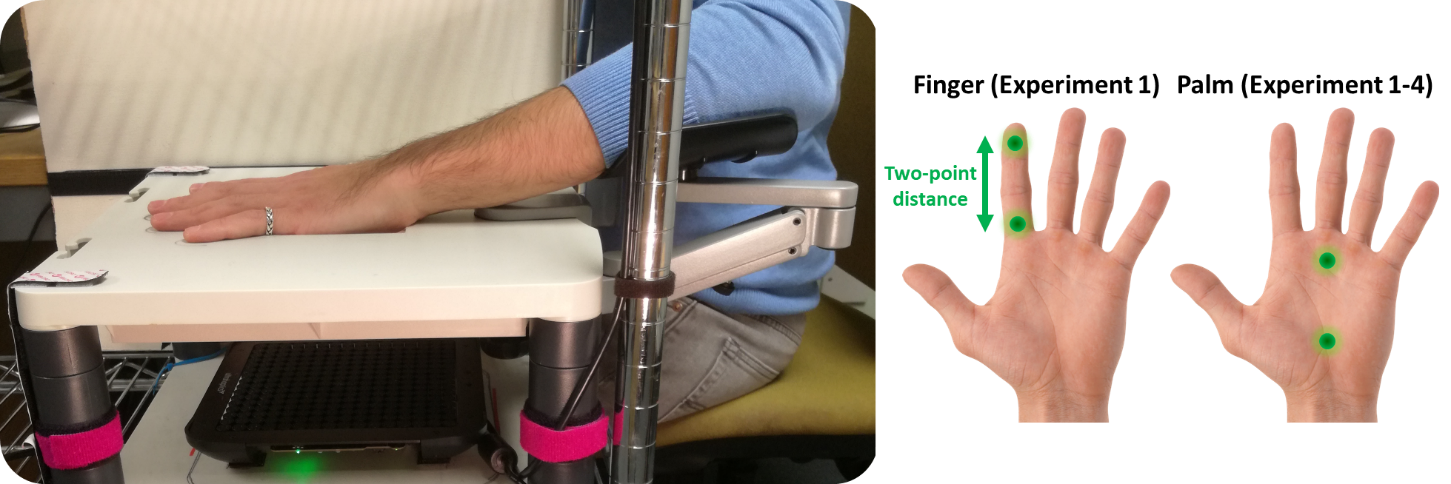

**Figure S1. Experimental Setup for Experiments 1-3.** The participants sat at a desk, resting their left elbow on an articulated armrest support and their left hand palm down on a plastic support, in correspondence of a 10 x 10 cm aperture. The view of the left hand was blocked by a fixed vertical plywood screen throughout the entire experiment. Airborne ultrasound stimulation was delivered through a STRATOS^TM^ Explore device located 12 cm below the participant’s hand. In Experiment 1, tactile stimuli were delivered either on the index finger or the palm of the hand. Given that Experiment 1 did not find a significant effect of the location of stimulation in Experiments 2-4 stimuli were delivered to the palm only.

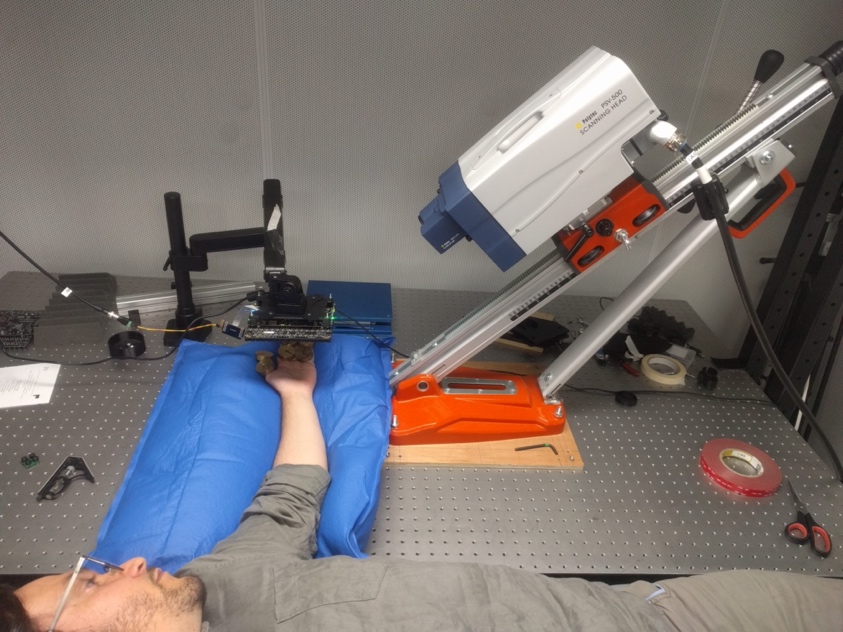

**Figure S2. Setup for the Laser Doppler Vibrometry in Experiment 4.** We used a Vibrometer to measure the indentation produced by the ultrasound stimulation on the participants’ hands. The LDV head was aimed at the participant hand at an angle of 45 degrees. The participants laid flat on a dentist chair, resting their arm on an optical table, supported by a vacuum cushion. The palm of the participants’ left hand faced upwards and received the vibrotactile stimulation from an ultrasound device suspended 12 cm above the hand and facing downwards. Skin indentation measures were taken for ultrasound stimuli of 50 Hz, 200 Hz, and perceptually-matched-200 Hz at the participant’s 2PD threshold and at suprathreshold (55 mm) distances.

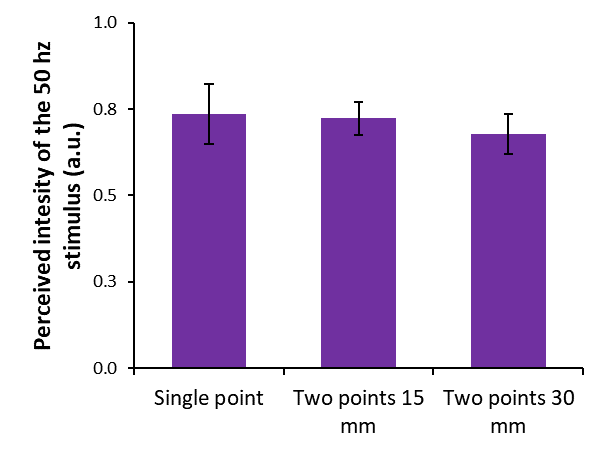

**Figure S3. Pilot Experiment on the Intensity Matching Task.** Prior to the beginning of Experiment 3 we conducted a small pilot study (n = 8) aiming to test whether using a single or a double point at 15 mm or 30 mm in the intensity-matching task produced statistically different results. Participants performed three blocks, one for each stimulation condition (single point, 15 mm, and 30 mm). Each block consisted in the staircase procedure described in the main text (see Methods) for the intensity matching task. A one-way repeated measures ANOVA showed that the differences between the three stimulation conditions were not significant (*F_2,14_* = 1.05; *p* = .375; η_p_^2^ = .131), suggesting that the participants’ performance in our intensity matching task was not affected by the number of points delivered or the distance between them.

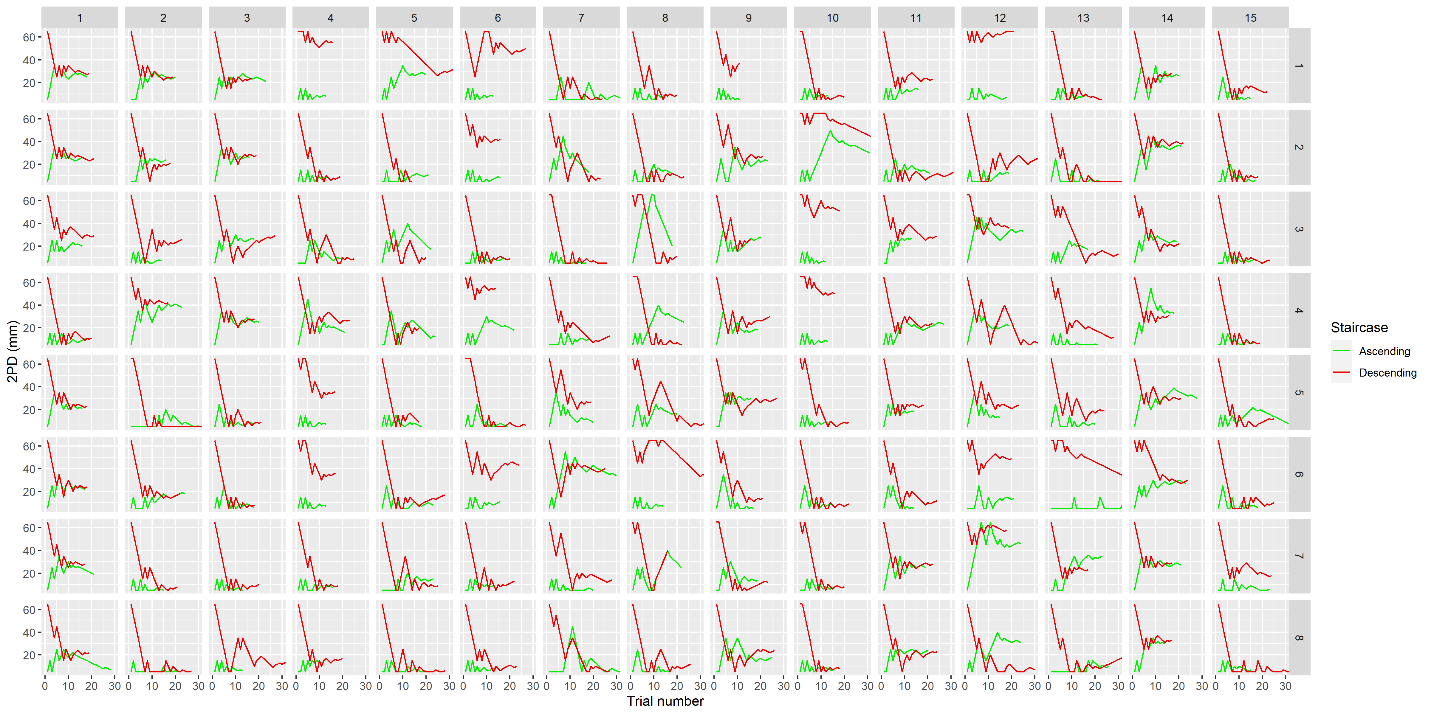

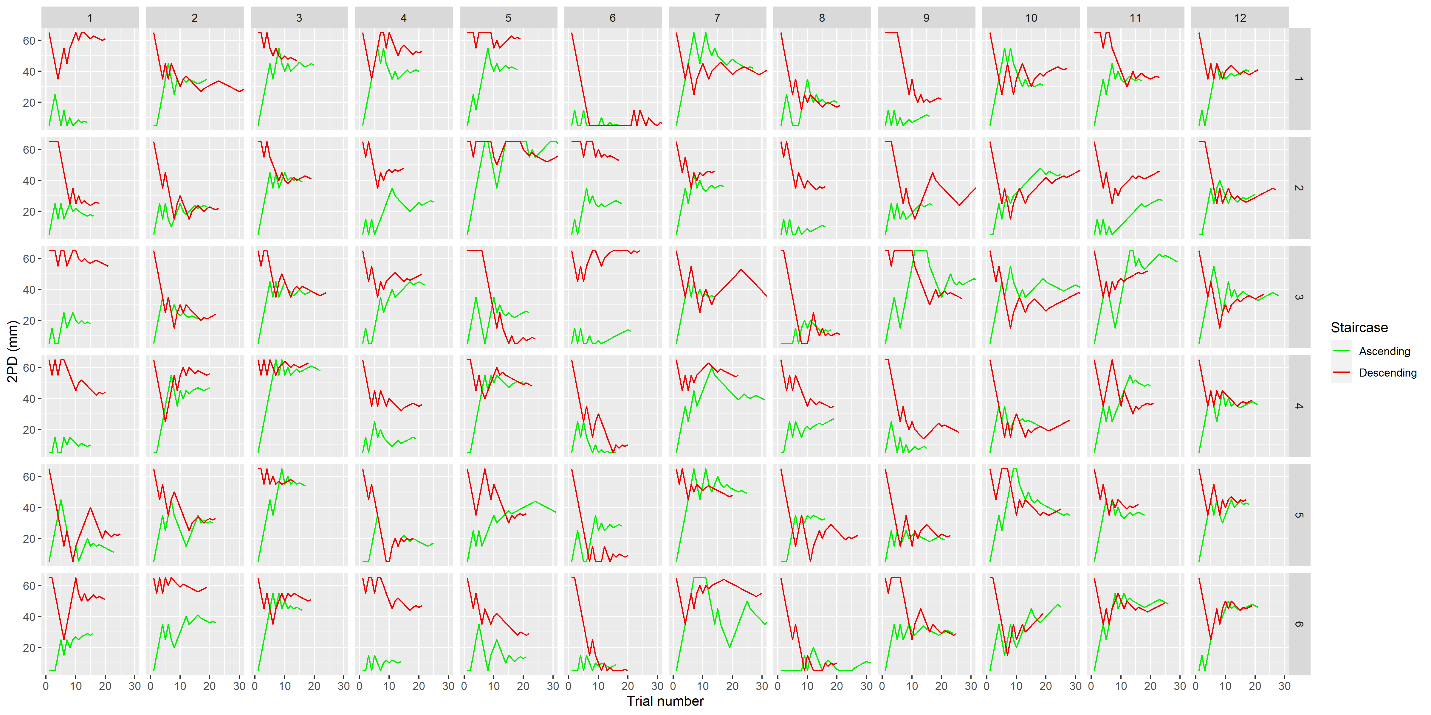

**Figure S4. Individual staircase data for each participant in each condition of Experiment 1 (top) and 2 (bottom).** Each plot represents the individual 2PD data for each participant (columns) and each block (rows). Red and green lines show participants responses to descending and ascending staircases, respectively. The participants’ 2PD threshold is calculated as the average of the last eight trials from both staircases in a block. Although we preregistered convergence between ascending and descending staircases as an exclusion criterium before data analyses, many more participants than expected showed poor convergence. This is most likely due to the relatively weak intensity of the ultrasound stimulus. As a consequence, we decided to deviate from our preregistration by also including data from blocks with poor convergence in our data analysis. We reasoned that a lack of convergence ultimately represents the difficulty of the participants in making the 2PD judgement in specific conditions. Moreover, it is important to note that a more liberal exclusion criterium only counts against our hypotheses by increasing the interindividual noise and lowering our statistical power. Thus, while using a more conservative staircase convergence criterium could potentially lead to a stronger effect than the one we found here, it would involve excluding a large number of participants, thus strongly limiting the generisability of our results. Fore example, applying the preregistered criterium of poor convergence in Experiment 1 would lead to the exclusion of seven participants out of 15 (i.e., 46.67% of the sample), an exceedingly high number. Importantly, applying such a strict exclusion criterium does not produce any change in the statistical inference of that experiment.
